# Simulated larvae dispersion of the invasive sun-coral (*Tubastrea* spp.) along Rio de Janeiro’s coast: the role of submesoscale filaments on offshore transport and connectivity

**DOI:** 10.1101/2024.10.23.619824

**Authors:** Leandro Calado, Bernardo Cosenza, Francisco Moraes, Caique Dias Luko, Damián Mizrahi, Fabio C. Xavier, Daniela Batista, Roberto Domingos, Sávio Calazans, Fernanda Araújo, Ricardo Coutinho

## Abstract

The spread of invasive species in marine ecosystems is a growing global concern, particularly in regions with high economic and ecological importance. sun corals (*Tubastraea* spp.) are scleractinians native to the Pacific Ocean that have spread along most of the Brazilian coast. This exotic species initially established populations in Rio de Janeiro, reaching high levels of abundance. Although the ecological aspects and impacts caused by this organism have been studied in detail, the natural mechanisms that drive its dispersal have attracted little attention. In this research, we focus on the offshore transport of sun coral larvae between Cabo de São Tomé and Ilha Grande Bay, RJ, investigating how submesoscale oceanographic features such as filaments, eddies and upwelling influence connectivity among different population. High-resolution numerical simulations were used to model the coastal dynamics, incorporating the influence of the Brazil Current, wind-driven circulation, and submesoscale structures. Larval dispersal was examined under different wind scenarios, including northeasterly winds that drive southward currents which enhance offshore transport via submesoscale filaments. Results show that submesoscale features, particularly filaments emerged from upwelling regions, play a significant role on sun coral larvae dispersion. These features act as pathways that connect larvae from coastal to offshore oil exploration areas, highlighting the importance of both natural and anthropogenic processes for the dissemination of this invasive species. This research provides critical insights into the ecological mechanisms governing the spread of invasive marine species, emphasizing the need for integrated coastal management strategies. Understanding how physical processes drive larval transport is essential for developing targeted control measures to mitigate the impact of invasive species like sun coral on native ecosystems and local economies. Furthermore, the study underscores the importance of monitoring both natural and anthropogenic influences on marine bioinvasions, particularly in regions with significant offshore industrial activities.

## Introduction

There is a growing concern regarding the spread of non-indigenous organisms in coastal regions. In Brazil, numerous studies assessed the impact that biological invasions have on the biodiversity of native flora and fauna, as well as on the economic activities associated with it [1], [2]. Particularly in the state of Rio de Janeiro, numerous exotic species have been the subject of extensive research due to their remarkable ability to colonize areas of economic or biodiversity importance. Among these species, the sun coral (*Tubastraea* spp.), has been a subject of increasing research over the past two decades [8]. The sun coral originates from the Pacific Ocean. It has initially spread throughout the Caribbean Sea in 1943, Gulf of Mexico in 1977 and the Southwestern Atlantic in the late 80’s, establishing population in the Campos Basin, off of Rio de Janeiro [3], [4]. These corals have been reported all along Rio’s coast, in areas such as Arraial do Cabo (AC), Ilha Grande Bay (IGB), Açú Harbor (AH) and Cagarras Island (CI), where populations have been established since the 1990s [4], [1], [5]. Once new populations are established, fertile adult colonies emit larvae - planulae - that are dispersed by ocean currents, since their swimming capacity is restricted at the microhabitat scale [6], [3], [7], [8], [5].

Although the coast of the state of Rio de Janeiro is densely populated by sun corals, the arrival of larvae from the open ocean is limited in this region [9]. Recently a comprehensive evaluation led by [10] focused on the distribution of sun coral larvae in the oceanic region of Cabo Frio. This study placed emphasis on their interaction with the Brazil Current (BC) and its influence by mesoscale dynamics. The findings from this study have illuminated that the BC can act as a dynamic barrier separating offshore and coastal areas. This discovery raises the hypothesis that the colonization of Sun Coral on the coast may be influenced by vectors responsible for transporting the larvae to regions in close proximity to the coastal continental shelf. This underscores the importance of understanding how coral dispersal mechanisms can be articulated or act synergistically.

The continental shelf in southeast Brazil can be categorized into outer, middle, and inner continental shelf, as detailed by [11]. From north of Cabo de São Tomé (CST) to Cabo Frio (CF) region, in contrast to the remainder of southeastern Brazil, there is a notable narrow continental shelf. Consequently, oceanic features like the BC with its meanders and eddies, exerts more influence on nutrient, plankton and physical properties changes between continental shelf and open ocean. South of Rio de Janeiro State, the continental shelf is larger and these open ocean processes are not dominant, however the frequency of incidence and intensity of cold front marked by the southwest wind are more common. As a result, a comprehensive understanding of the coastal region must account for the intricacies of the continental shelf itself, which are influenced by the wind patterns, mesoscale dynamics of the Brazil Current and submesoscale processes on the shelf. This coastal area has already witnessed conditional sun coral infestation, primarily in the Arraial do Cabo region. These circumstances are noteworthy due to their unique nature, as they appear to be stable despite their exotic origins. However, uncovering the potential primary sources of this colonization points towards connections with offshore oil and gas exploration along the southeastern coast of Brazil. This may arise from the support equipment that frequently docks in coastal ports or from the potential influx of larvae from exploration zones.

*Tubastraea* spp. can be effectively spread through offshore oil and gas operations, particularly via vessels and industrial structures. As noted by [8], *Tubastraea* spp. have shown a strong ability to survive on stationary, slow-moving structures such as oil platforms, support vessels, and equipment, especially those used in the oil and gas offshore industry. This observation implies that support vessels can play a major role as vectos of introduction on the introduction of *Tubastraea* spp., facilitating their spread along the Brazilian coast. According to [12], there is evidence that mechanically removing sun coral colonies can lead to significant larval releases during high fecundity periods. This on-site evidence underscores the need for cautious consideration when opting for mechanical removal as the preferred method to curb the proliferation of sun coral. These techniques may lead to additional risks of anthropogenic dissemination due to the regenerative capacity of coral fragments [13], [14] and the release response of large numbers of larvae as a result of the stress caused by mechanical removal [12]. This is a possible explanation of the high abundance of this exotic organism in IGB [15], [16], since mechanical sun coral removal protocols have historically been practiced in the IGB nautical center that supports offshore oil and gas activity [17].

The expansion of offshore oil and gas exploration areas may facilitate the oceanic dispersion of sun coral, once it leads to an increase in the availability of consolidated substrate on the oil and gas exploration areas . [10] showed that in the Brazil Current domain some larvae interact with Cabo Frio eddies, restricting their distribution but remaining at regions where the production fields are located, while others are captured by BC increasing their dispersal, however without reaching the coast. On the nearshore zone, [18] approached the coastal dispersion of sun coral larvae restricted to the Arraial do Cabo internal region, nonetheless dispersal and connectivity with other infested coastal regions has not been fully addressed. Furthermore, when it comes to coastal-offshore exports its not clear if the coastal populations can be a larval source to colonize offshore structures.

The evolution of the ocean dynamics on the continental shelf is influenced by the wind regime that acts on the shelf determining the direction of the coastal current [11]. While coastal upwelling stands as the primary phenomenon in these regions, persistent winds from the Northeast, occasionally combined with the inlet of cold fronts carrying Southwest winds, serve as the primary factors in generating upwelling or subsidence, and additionally dictates the direction of currents on the continental shelf. This change in currents due to wind regime, especially during periods of cold front entry, might drive the dispersion of larvae in the region. Furthermore, although the coastal region has its own dynamics, [19] and [20], show that the interaction among the eddies of the Brazil Current plays a fundamental role in coastal dynamics.

In addition to the influence of winds and BC meanders on the shelf dynamics, submesoscale phenomena also play a major role on the circulation. Submesoscale currents have typical length scales that range from 100s m-10s km [21], and are dynamically defined as currents with 𝒪(1) Rossby numbers (*Ro* = *ζ*/*f*, where *ζ* is relative vorticity and *f* is the planetary vorticity). Submesoscale phenomena can be triggered by flow-topography interactions, especially in separation regions where enhanced lateral shear against topography ejects filaments and eddies [22], [23], [24], [25], [26]. Nearshore lagrangian transport and connectivity are intimately linked to submesoscale currents, which aggregate material [27], [28] and open up transport pathways across the shelf [29].

Submesoscale filaments associated with the Cabo Frio upwelling system have been reported as common features observed in numerical simulations [30]. The observation of these submesoscale filaments has been a challenge due to there small spatial scales, and to SST biases in microwave-based satellite datasets [31]. However, in clear-sky conditions, these filaments can be seen using the recently-available infrared-radiometer data (Level 2 with 1-km resolution at nadir) from the Sentinel 3 satellite (Figure 1a). These cold submesoscale filaments, arising from Cabo Frio (23^°^S) and Cabo de São Tomé (22^°^S), are associated to strong chlorophyll concentrations (Figure 1b), and might be essential for the cross-shelf export of water properties, carbon and larvae in the region. In particular, the filaments must be key features for the regional dispersal of sun coral larvae since populations have been reported in Arraial do Cabo and Açu Harbor.

**Fig 1.**
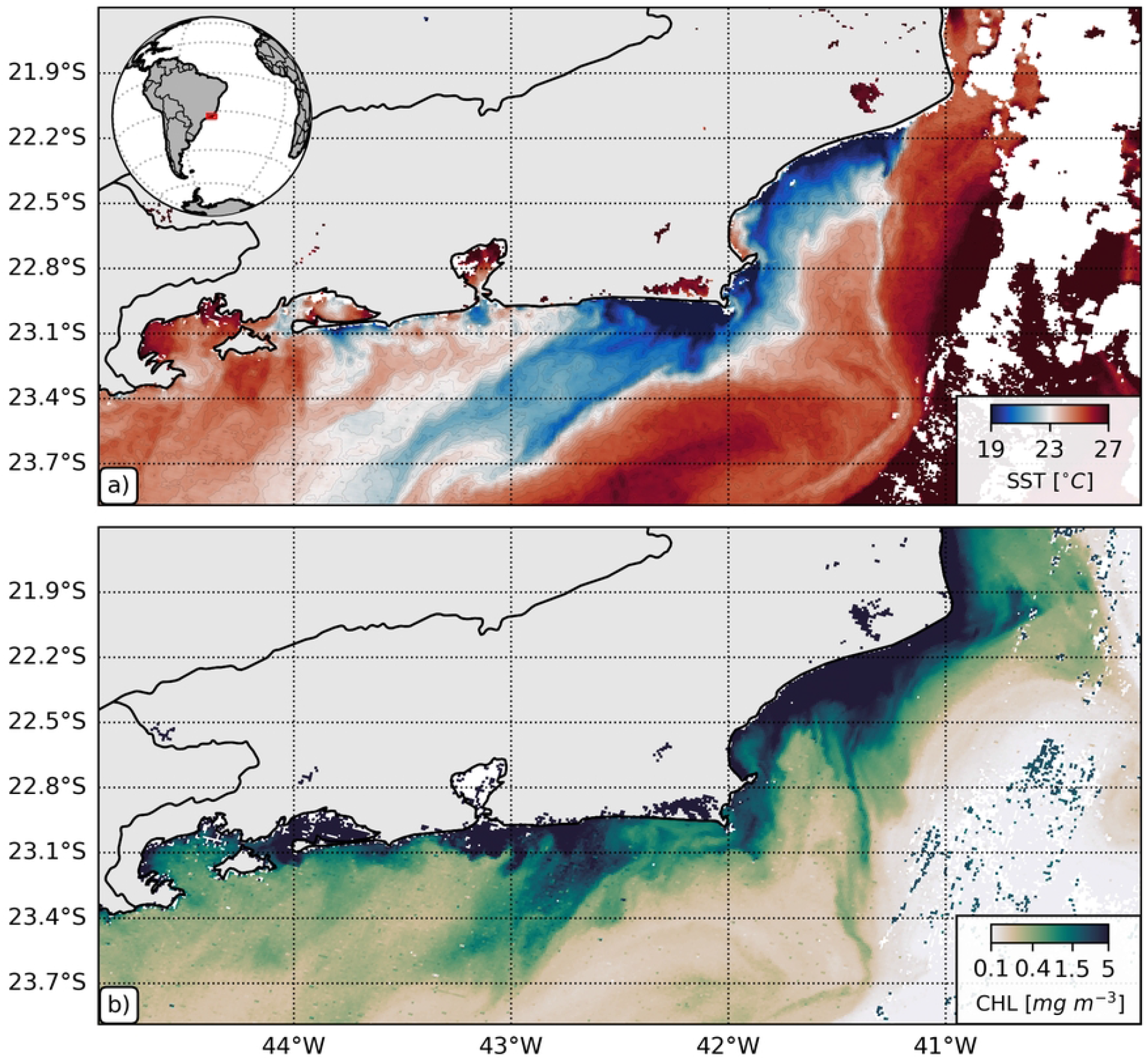
Submesoscale filaments along the coast of Rio de Janeiro (May 07 2024): a) sea surface temperature (SST); and b) chlorophyll concentration (CHL). Level 2 satellite images from Sentinel 3A.

In this study, we investigate the role of submesoscale phenomena in establishing transport pathways and connectivity of sun coral along the coast of Rio de Janeiro (Brazil). Our goal is to identify how the continental shelf circulation controls coastal inter-connectivity between colonized regions, and to test the offshore larvae exportation hypothesis. We focus our investigation in two realistic case studies where different wind conditions are applied. In these scenarios, different wind forcing modulate the submesoscale activity by changing the shelf circulation. In the course of this study, we conducted experiments, building upon the numerical implementation presented by [10], with the aim of investigating the interchangeability of sun coral larvae within the coastal region. This implementation involved the incorporation of a third coastal grid with an approximate resolution of 600 m. On Section 2, we present the numerical model we implemented, while on Section 3, we detail our larval dispersal experiments. On Section 4, we present and discuss our Results, and on Section 5 we present our Conclusions.

## Material and Methods

### Model Implementation

The study area includes the region between Cabo de São Tomé (21.9^°^*S*) and Ilha Grande Bay (23.4^°^*S*; Figure 2). The main goal is to examine how larval dispersion occurs from source points such as rocky shores, ports, and platforms associated with offshore oil and gas activities. These anthropogenic structures act as potential vectors for the dispersion of sun coral larvae when vessels transit between them and the near coast, particularly in ports. Figure 2 highlights source locals in the Southeast where larval emission is likely to occur, including four coastal release sites: Arraial do Cabo (AC), Ilha Grande Bay (IGB), Açú Harbor (AH) and Cagarras Island (CI). Açú Harbor was chosen as a source point due to its operation by vessels from the oil and gas industry, and reports of significant sun coral (*Tubastraea* spp.) presence in both natural rocky environments and artificial substrates. Offshore oil fields are marked in white with green contours, and two key points, P1 and P2, in the Campos Basin are identified as potential larval arrival sites.

**Fig 2.**
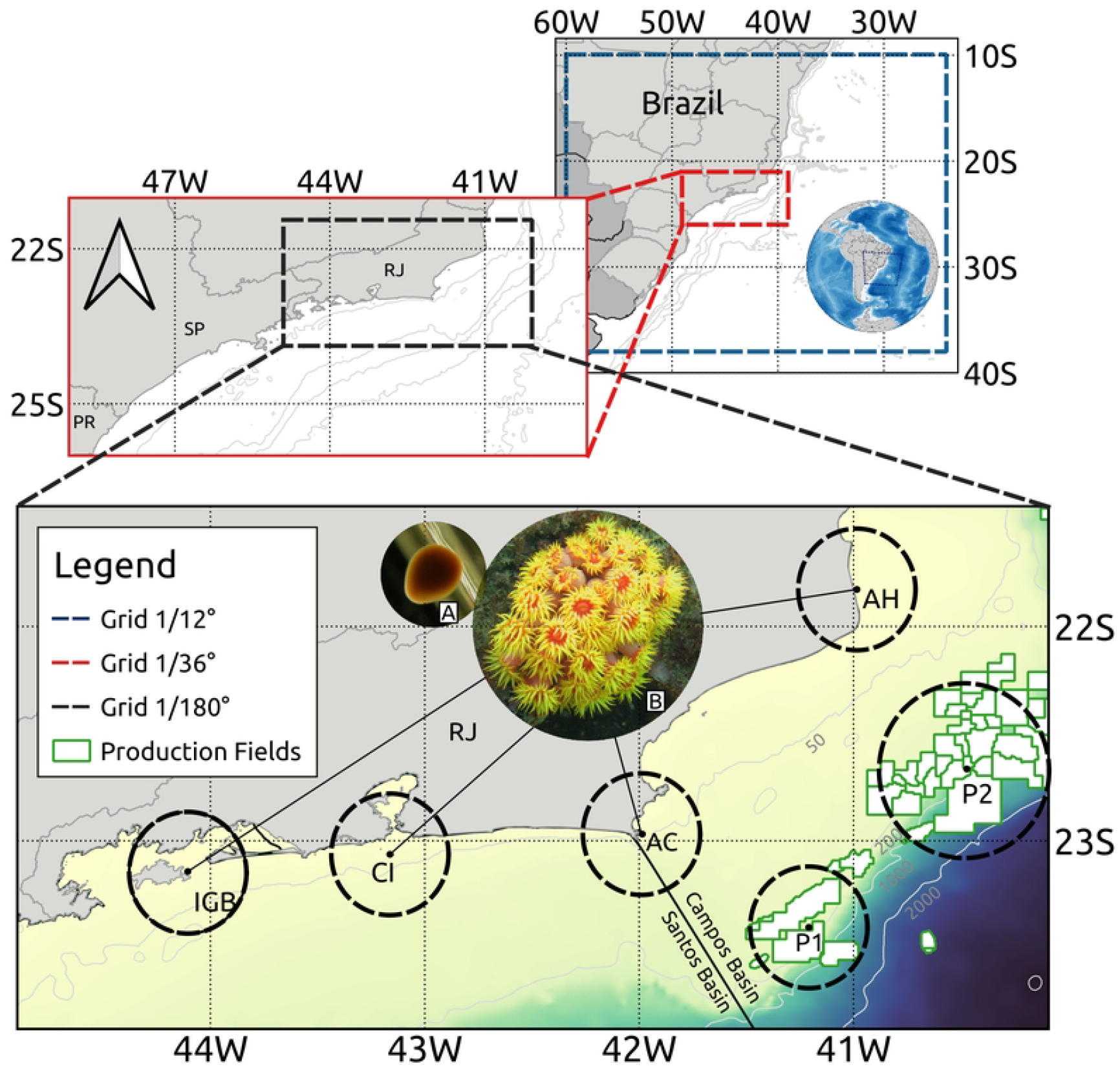
Model grids representing the study area. Coarse resolution grids used for nesting and on the higher one, it is highlighting the four locations of larvae release - Ilha Grande Bay (IGB), Cagarras Island (CI), Arraial do Cabo (AC) and Açú Harbor (AH). The offshore oil production fields are white shaded with green contour, the gray lines are the isobaths. Panel A: photo of the sun coral larva (pyriform shaped) settling and, panel B: adult colony open potential for larval emission).

The Regional Ocean Model System (ROMS) serves as the primary model employed in this study [32]. ROMS is a three-dimensional regional model that employs the primitive equations of motion in terrain-following vertical coordinates. ROMS is a modular framework that accommodates the inclusion of atmospheric and tidal components, as well as integration with other global models at its open boundaries. Our implementation follows the progression established by [10]. Here, we used three nested grids, outlined in Figure 2. The model is forced using atmospheric forcing from ERA5 [33], oceanic forcing at the boundaries from GLORYS12V1 [34], and astronomical tide from the TPXO global tide model [35], [36].

The submesoscale phenomena present in the Cabo Frio region, located off the southeastern coast of Brazil, resolved nesting three levels of resolution: a parent grid at 1/12^°^, a child grid at 1/36^°^, and a Gchild grid at 1/180^°^(Figure 2). This hierarchical approach allowed us to capture the intricate details of eddies and filaments with varying degrees of precision. Through this gradual reduction in horizontal resolution, we reach a grid spacing of 600 m within the Cabo Frio region (Gchild domain). This progressive refinement offers unparalleled insights into the submesoscale features prevalent in the region.

The validation of the hydrodynamic model is detailed in [10] and using real-world observational data, such as ARGO profilers measurements (Figure S1 in the Supplementary Material) for the Gchild grid. The strong agreement between observed and simulated temperature-salinity structures highlights the model’s ability to accurately replicate the complex dynamics of the Brazil Current system, reinforcing its potential for broader ecological and environmental applications.

The model incorporates Lagrangian derivatives characterized by the traits commonly associated with lecithotrophic larvae. These larvae have reduced swimming abilities, and feed on energy reserves stored in internal vesicles, exhibiting a density profile akin to that of sun coral larvae [37], [38]. Due to this reduced swimming ability, we simulate the sun coral larvae as depth-keeping organisms [39] and only advect them with horizontal velocities [40], [41]. The larvae are released in proximity to the water’s surface and their dispersion is computed at each ROMS time step.

It has been shown for several species of corals from different regions that, the larval competency period appears to be the primary factor influencing their spatial distribution [37]. The competency window is the lapse of time during which the larvae can settle on a substrate and successfully undergo metamorphosis into early juvenile recruits [42]. As these larvae do not feed on plankton, this period is constrained by the nutritional reserves allocated inside internal vesicles during embryogenesis [43]. Several studies have linked the larval period of competence window to the dispersal potential of organisms that emit non-planktonic larvae [44], [45], [46]. sun coral larvae can spend up to 90 days in the water column before recruitment happens [13], [47], [9], which contributes to the spread success of this species. Here, our study cases have the objective of understanding the role of submesoscale dynamics on larvae dispersal. Thus, since the time-scales of these processes can range from days to weeks, we limit our case study experiments to 20 days. We show that this is enough time for submesoscale currents to affect inter-connectivity of different populations and to promote offshore export to oil and gas exploration regions.

## Experiments

Two experiments were conducted with the aim of comprehending potential source areas of sun coral larvae and areas suitable for colonization, based on the coastal hydrodynamics. We consider scenarios involving different wind conditions, and the combined influence of tidal currents, which modulate the progression of coastal currents and submesoscale filaments originated in theses scenarios. The prevailing wind patterns in the region are the primary drivers of these currents, as explained by [11] and [48]. Thus, scenarios with different wind regimes are simulated to investigate how sun coral population in AC, AH, CI and IGB (Figure 2) are interconnected among each other and with offshore oil platforms.

The frequent change in wind conditions in this region is due to the passage of atmospheric cold fronts that disturb the prevalent northeastward winds, associated to the South Atlantic High, to intermittent southwestward winds. We design our study case experiments to capture these two scenarios, dividing them into: 1) Prevailing Northeast Winds Scenario; and 2) Cold Front Winds Scenario (Figure 3). Both experiments were carried out for 20 days, focusing on how the fast submesoscale processes can modulate the initial dispersal of sun coral larvae.

**Fig 3.**
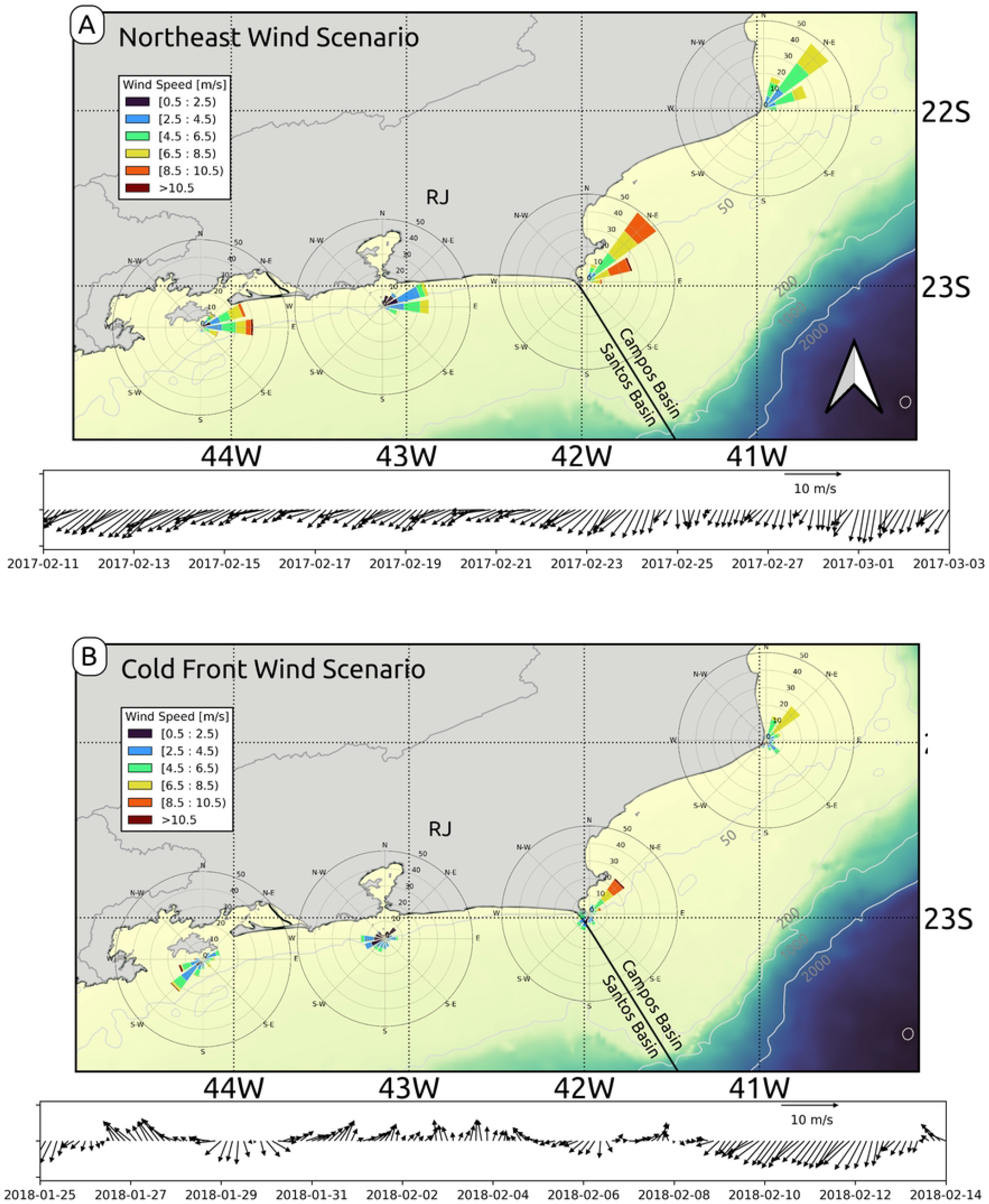
Wind patterns for two scenarios: (a) Northeast wind scenario and (b) Transition from northeast to cold front winds.

Figure 3 presents the wind roses at four key dispersal points of the invasive sun coral. Each panel corresponds to a specific scenario, displaying an average stick plot of the wind field over the high-resolution grid for the designated simulation periods. These stick plots provide a detailed view of the wind direction and speed, allowing for a comparative analysis of the meteorological conditions that shape larval dispersal patterns in the study area.

Figure 3a and b show distinct differences between the two proposed wind scenarios on the model simulations. Figure 3a captures the dominant northeast wind pattern, which is typical of the southeastern coast of Brazil. This northeast wind regime is associated with stable atmospheric conditions and a steady influence on surface currents. The consistent northeast wind plays a significant role in shaping nearshore hydrodynamics, fostering upwelling events, and influencing the local ocean circulation. These dynamics are crucial for understanding how larvae of *Tubastraea* spp. are transported along the coast, as this wind-driven circulation creates opportunities for the larvae to remain in coastal waters or be exported offshore, aiding in their spread.

In contrast, Figure 3b illustrates a shift in wind patterns due to the arrival of a cold front, a common meteorological phenomenon in the region. As the cold front arrives, the winds turn to the southwest, significantly altering the regional wind regime. This shift is critical for larval dispersal because it can disrupt the established northeast wind-driven currents and lead to a temporary reversal in surface flows. As the southwest cold front winds increase in intensity, they can generate new circulation features, such as downwelling near the coast or changes in the strength and direction of alongshore currents. The alternation between northeast and cold front wind events observed in this scenario reflects the real atmospheric variability experienced in this region. The transition between these wind patterns underscores the complexity of coastal weather systems and their profound impact on ocean circulation, which, in turn, affects biological processes like larval transport.

## Results and Discussion

### Submesoscale regimes under different wind conditions

The presence of submesoscale features off Rio de Janeiro’s coast [30] holds the potential to facilitate the export of coastal larvae to offshore areas. Submesoscale features include eddies, fronts and filaments which develop on temporal and spatial scales between the mesoscale and turbulence (typical length-scales from 100s m - 10s km), and are usually characterized by 𝒪(1) Rossby numbers (*ζ/f*). These phenomena play a crucial role in redistributing physical and biogeochemical properties [49], such as heat, nutrients, and organisms, thereby linking broader coastal and oceanic dynamics and significantly influencing marine circulation and dispersal mechanisms.

Submesoscale motions can be triggered by flow-topography interactions [25], [26] which, on the shelf, can be modulated by the wind-driven circulation that alters how currents interact with topography. Our experiments focus on two scenarios that represent the main wind forcing observed in the region: 1) The Northeast Wind Scenario, and 2) The Cold Front Wind Scenario (Figure 3).

Under Northeast winds, the coastal circulation is characterized across the whole shelf by southwestward flowing currents. In this scenario, several filaments form and transport cold upwelling water southwestward (Figures 4a-c and 5a). The most prominent filament originates at Cabo Frio (42^°^*W*), although other filaments are also observed along the coast, e.g.: i) Cabo de São Tomé (41^°^*W*); ii) west of Baía de Guanabara (43.5^°^*W*); iii) Ilha Grande Bay (44.1^°^*W*); and iv) Ponta da Juatinga (44.5^°^*W*).

**Fig 4.**
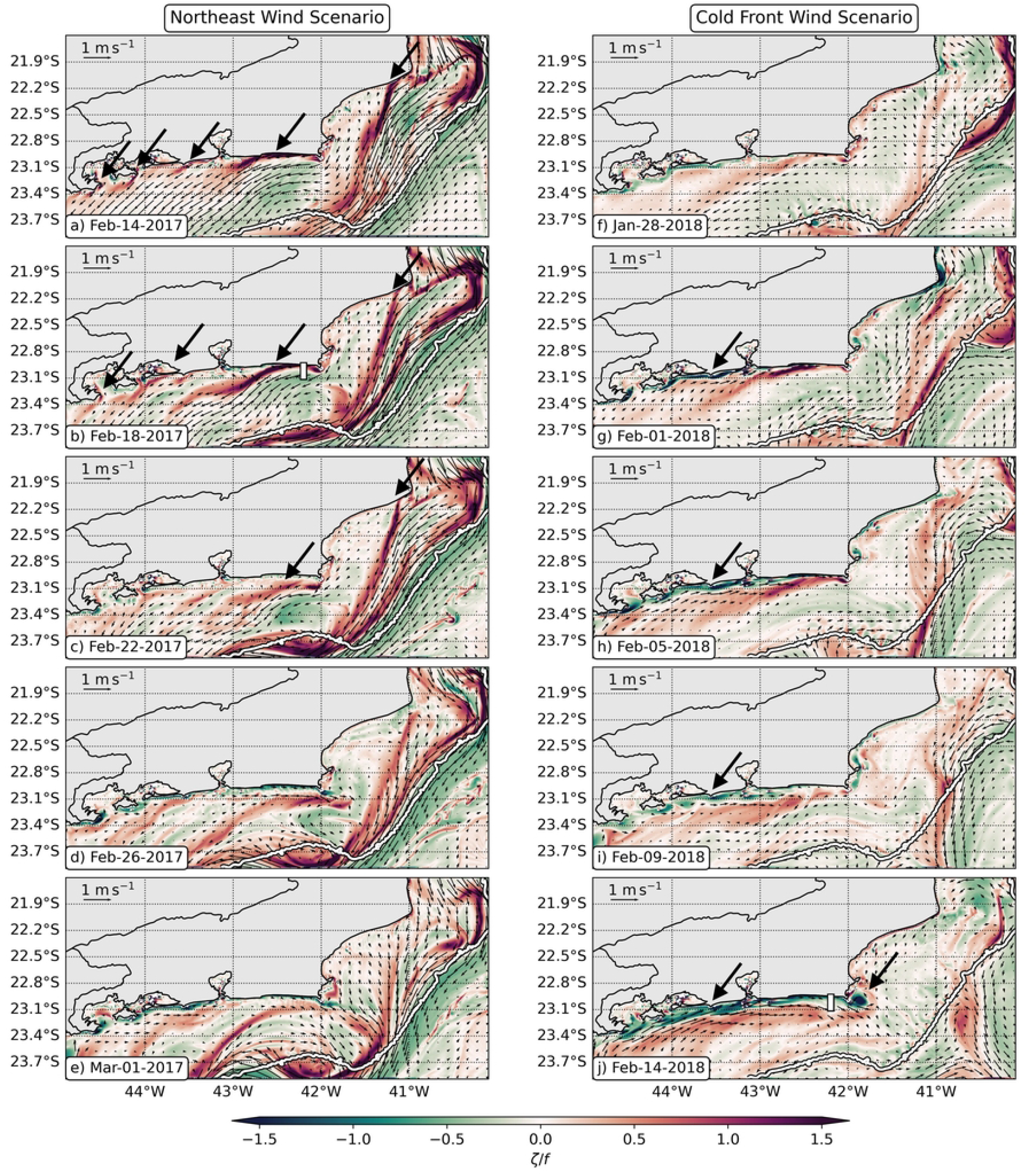
Submesoscale filaments and eddies off Rio de Janeiro’s coast for the Northeast wind (left column) and Cold Front wind (right column) cases. Horizontal maps of surface Rossby number (*ζ*/*f*) on: a) Feb 14 2017, b) Feb 18 2017, c) Feb 22 2017, d) Feb 26 2017, e) Mar 01 2017, f) Jan 28 2018, g) Feb 01 2018, h) Feb 05 2018, i) Feb 09 2018, and j) Feb 14 2018. The white bars on panels b and j show the position of the vertical sections displayed on Figure 5. The white contours on all panels show the 200 m isobath, and the big dark arrows highlight the cyclonic (red) and anticyclonic (green) submesoscale filaments.

These filaments are generated on separation regions [22], [50], where the coastal current interacts with topography increasing lateral shear and vorticity (Figures 4a-c and 5e). Intense cyclonic vorticity is ejected from CF, giving rise to the submesoscale filaments (the narrow regions with 𝒪(1) *ζ/f* ; Figure 4e), where Lagrangian material typically accumulates due to enhanced strain rates (Figure 4c). Around February 26 2017, the northeast winds weakened on the west part of the domain leading to less southwestward currents (Figure 4d). Consequently, submesoscale cyclonic filaments no longer dominate the shelf circulation west of Cabo Frio (42^°^W). In fact, by March 1, we observe that a mesoscale cyclone forms in the Cabo frio region. This mesoscale cyclone brings southeastward currents to Arraial do Cabo and is another potential driver of larvae dispersal.

The presence of submesoscale features off Rio de Janeiro’s coast [30] holds the potential to facilitate the export of coastal larvae to offshore areas. Submesoscale features include eddies, fronts and filaments which develop on temporal and spatial scales between the mesoscale and turbulence (typical length-scales from 100s m - 10s km), and are usually characterized by 𝒪(1) Rossby numbers (*ζ/f*). These phenomena play a crucial role in redistributing physical and biogeochemical properties [49], such as heat, nutrients, and organisms, thereby linking broader coastal and oceanic dynamics and significantly influencing marine circulation and dispersal mechanisms.

Submesoscale motions can be triggered by flow-topography interactions [25], [26] which, on the shelf, can be modulated by the wind-driven circulation that alters how currents interact with topography. Our experiments focus on two scenarios that represent the main wind forcing observed in the region: 1) The Northeast Wind Scenario, and 2) The Cold Front Wind Scenario (Figure 3).

Under Northeast winds, the coastal circulation is characterized across the whole shelf by southwestward flowing currents. In this scenario, several filaments form and transport cold upwelling water southwestward (Figures 4a-c and 5a). The most prominent filament originates at Cabo Frio (42^°^*W*), although other filaments are also observed along the coast, e.g.: i) Cabo de São Tomé (41^°^*W*); ii) west of Baía de Guanabara (43.5^°^*W*); iii) Ilha Grande Bay (44.1^°^*W*); and iv) Ponta da Juatinga (44.5^°^*W*).

These filaments are generated on separation regions [22], [50], where the coastal current interacts with topography increasing lateral shear and vorticity (Figures 4a-c and 5e). Intense cyclonic vorticity is ejected from CF, giving rise to the submesoscale filaments (the narrow regions with 𝒪(1) *ζ/f* ; Figure 4e), where Lagrangian material typically accumulates due to enhanced strain rates (Figure 4c). Around February 26 2017, the northeast winds weakened on the west part of the domain leading to less southwestward currents (Figure 4d). Consequently, submesoscale cyclonic filaments no longer dominate the shelf circulation west of Cabo Frio (42^°^W). In fact, by March 1, we observe that a mesoscale cyclone forms in the Cabo frio region. This mesoscale cyclone brings southeastward currents to Arraial do Cabo and is another potential driver of larvae dispersal.

In the Cold Front Wind Scenario, the simulation starts with weak northeast winds (Figure 3b) and we observe weak cyclonic filaments on the shelf (Figure 4a). Around February 1 2018, a cold atmospheric front enters the domain and the winds shift from northeast to southwest winds. This wind shift leads to northeastward currents that form narrow anticyclonic submesoscale filaments that remain trapped near the coast (Figure 4g). From February 5 to February 8, the winds shift to northeast winds again (Figure 3b), and some southwestward cyclonic filaments are ejected into the offshore region (Figure 4h) from Cabo de São Tomé (41^°^*W*) and Cabo Frio (42^°^*W*).

This fluctuating pattern reveals that when Cold Atmospheric Fronts arrive in the region, the anticyclonic filaments might act to trap the larvae near the coast. This is confirmed by the stronger normalized strain rates near the coast (Figure 5d), that restrict the accumulation of larvae to the nearshore region. However, the shift to northeast winds can still allow some of those larvae to escape due to the formation of the southwestward cyclonic filaments.

**Fig 5.**
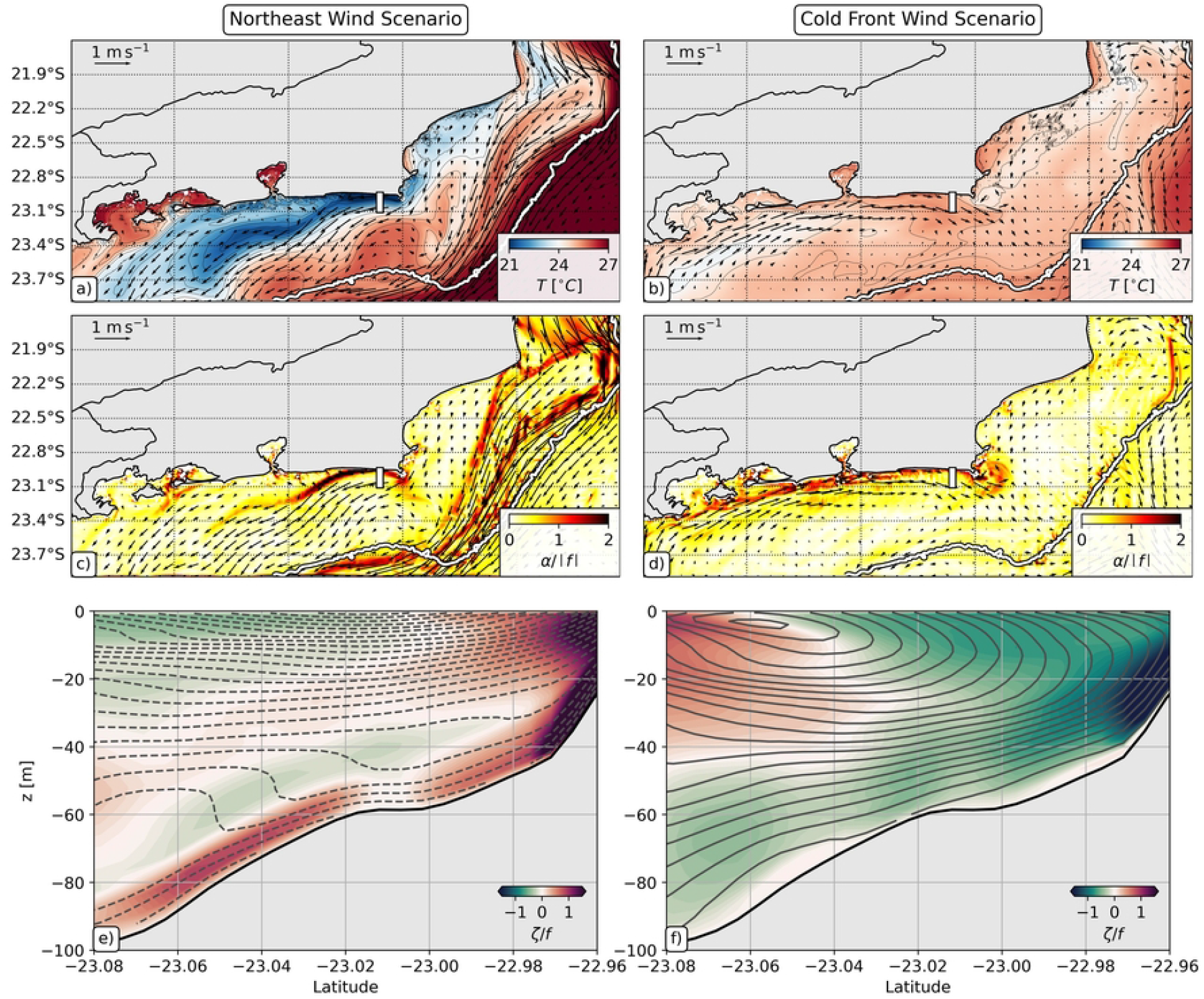
Submesoscale filaments for the NE wind case scenario on Feb 18 2017 (left column) and for the Cold Front wind scenario on Feb 14 2018 (right column). Horizontal maps of: (a-b) sea surface temperature, and (c-d) normalized strain rate (*α*/ | *f* |). The white contours on panels a-d show the 200 m isobath. Vertical sections of: (e-f) Rossby number (*ζ*/*f*). The position of the vertical sections displayed on panels e and f is shown as white bars on panels a-d and on Figure 4b,j. The solid (dashed) isolines on panels e and f represent lines of constant eastward (westward) velocities.

Eventually, if the Cold Front Winds are persistent, the northeastward currents can also eject submesoscale features offshore. In this scenario, the lateral shear against topography amplifies anticyclonic vorticity (Figure 4j and Figure 5f), and we observe the formation of submesoscale anticyclonic features downstream of Cabo Frio (Figure 5f, similarly to [51]). Even though this was not observed in our study case, these anticyclones could trap particles and travel offshore, also contributing to larvae export.

To understand how these submesoscale filaments and eddies affect the dispersion of sun coral larvae on this region, we analyze the trajectories of simulated Lagrangian particles in each experiment. We focus on the effect of horizontal transport, but we also acknowledge that the different wind scenarios and submesoscale regimes can also affect temperature (Figures 5a-b) and, thus, larvae dispersion might also be affected due to changes on environmental conditions. The results of our larvae dispersion experiments are shown on the next section.

### Larval Dispersion Along Rio de Janeiro’s Coast

The larvae dispersion experiments show that larvae transport are mainly driven by the submesoscale regimes modulated by the wind-driven circulation. We observe that different regimes affect patterns of offshore transport, near coast entrapment and, consequently, connectivity between different colonies.

Under Northeast wind conditions, the dispersion patterns along the shelf revealed that larvae tend to move southwestward, with significant aggregation in narrow filaments (Figures 6a-c). The aggregation of larvae is often co-located with regions of 𝒪(1) Rossby number, confirming that submesoscale filaments play a major role on opening transport pathways for larvae dispersal across the shelf [29, e.g.,]. The aggregation of material on filaments is a typical feature of submesoscale currents [27], [28], [49] due to the strong strain rates associated to these processes (e.g., Figure 5c).

**Fig 6.**
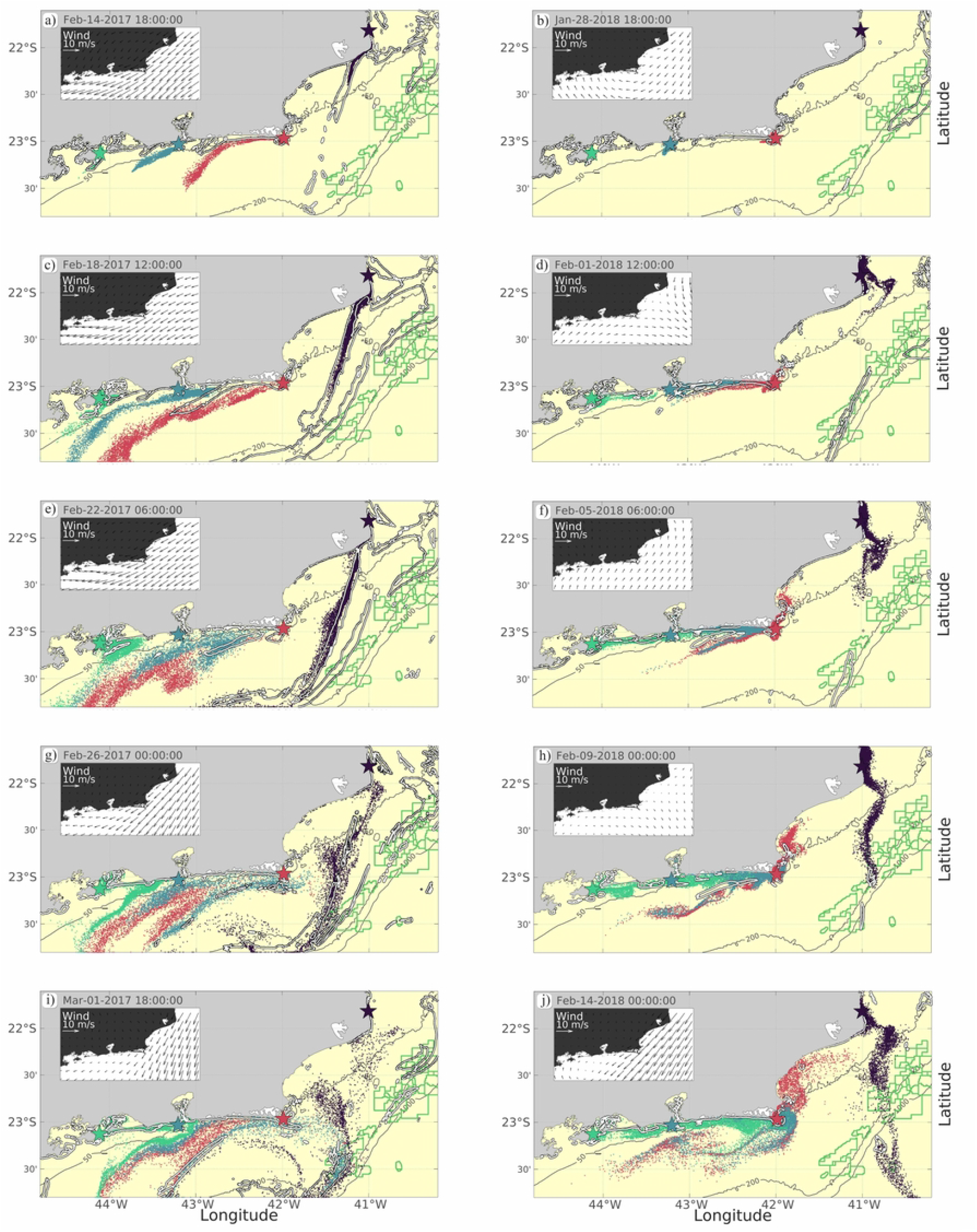
Larval dispersal simulation for each dispersal point in the wind direction scenario (northeast wind and cold front wind). The colors of the derivatives indicate their region of origin. The white lines correspond to the Rossby number O(1).

This scenario favors the offshore transport of larvae besides supporting southwestward transport from AC to CI and from CI to IGB (Figures 6a,c,e and 7c,e). Here, we observe that larvae can be transported from the coast towards oil platforms as exemplified by the drifters from AH that reach P1 (Figures 6e,g and 7a). After the northeast winds weaken at the western portion of the domain, northeastward currents contribute to advect IGB larvae to CI (Figures 6i and 7g) and, moreover, the formation of a mesoscale cyclone off Cabo Frio supports larvae transport from CI to AC, and from AC to P1 (Figures 6a and 7b).

In the scenario characterized by a shift to Cold Front winds, a markedly different dispersal pattern emerges. The southwest wind regime tends to concentrate larvae closer to the coast and lead to northeastward transport (Figures 6b,d,f), altering the dispersal pathways observed under northeast winds. In this scenario, larvae originating from IGB, in Sepetiba region and the Cagarras Island, near Guanabara Bay, exhibit a strong tendency to remain within the coastal zone (Figures 6d,f,h,j and 7h). This retention is likely driven by the reversal of surface currents, which now favor an onshore movement, trapping larvae in nearshore environments. However, this scenario is characterized by fluctuating conditions due to the alternation between cold front winds and the prevalent northeast winds. Thus, some larvae can still escape the near-coast entrapment and be transported offshore by cyclonic filaments (Figures 6f,h,j and 7d,f,h). Consequently, even in this scenario coastal larvae from AH can reach oil exploration regions (Figures 6h,j and 7b).

The concentration of larvae in coastal areas, especially within embayments like Sepetiba and Guanabara Bay observed in the Cold Front Winds scenario, is likely influenced by a combination of wind-driven currents and the local tidal regime. The tidal forces in these areas can further promote the retention of larvae, limiting their dispersion to offshore regions. This is particularly evident in Figure 6, panels g and h, where the southwest wind scenario results in a higher percentage of larvae being retained near the coast. The dynamics within these embayments can have profound implications for local ecosystems, as the introduction of invasive *Tubastraea* spp. species may alter the ecological balance within these relatively enclosed environments. The presence of these larvae in coastal waters may also increase the likelihood of colonization in nearshore natural rocky environments, artificial structures, such as harbors, marinas, and other man-made infrastructures.

Figure 8 (panels a and b) presents connectivity matrices depicting the relationship between the source and destination sites for larval dispersal under two distinct wind scenarios. These matrices provide a quantitative analysis of the dispersion patterns and the concentration of larvae at various destination sites, reinforcing the earlier observations about how wind regimes influence larval transport. The percentage values in these matrices highlight the extent of connectivity between source regions and the final settlement areas for larvae, offering a deeper understanding of the dynamics at play in each scenario.

**Fig 7.**
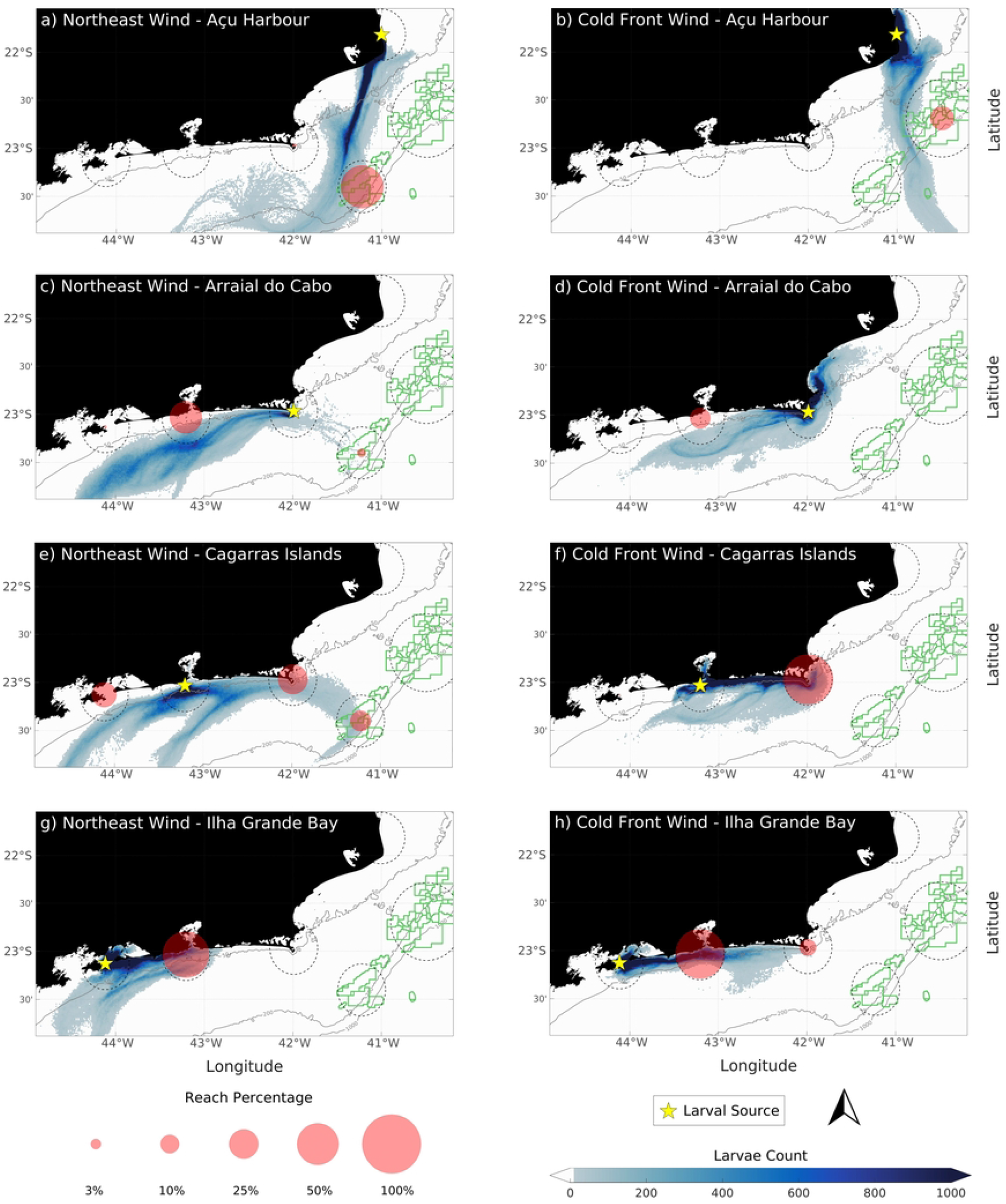
Integrated density plume of larval dispersion (blue plume) for each wind direction (Northeast wind and cold front wind). The pink circles indicate the final population of larvae at the end of the simulation.

**Fig 8.**
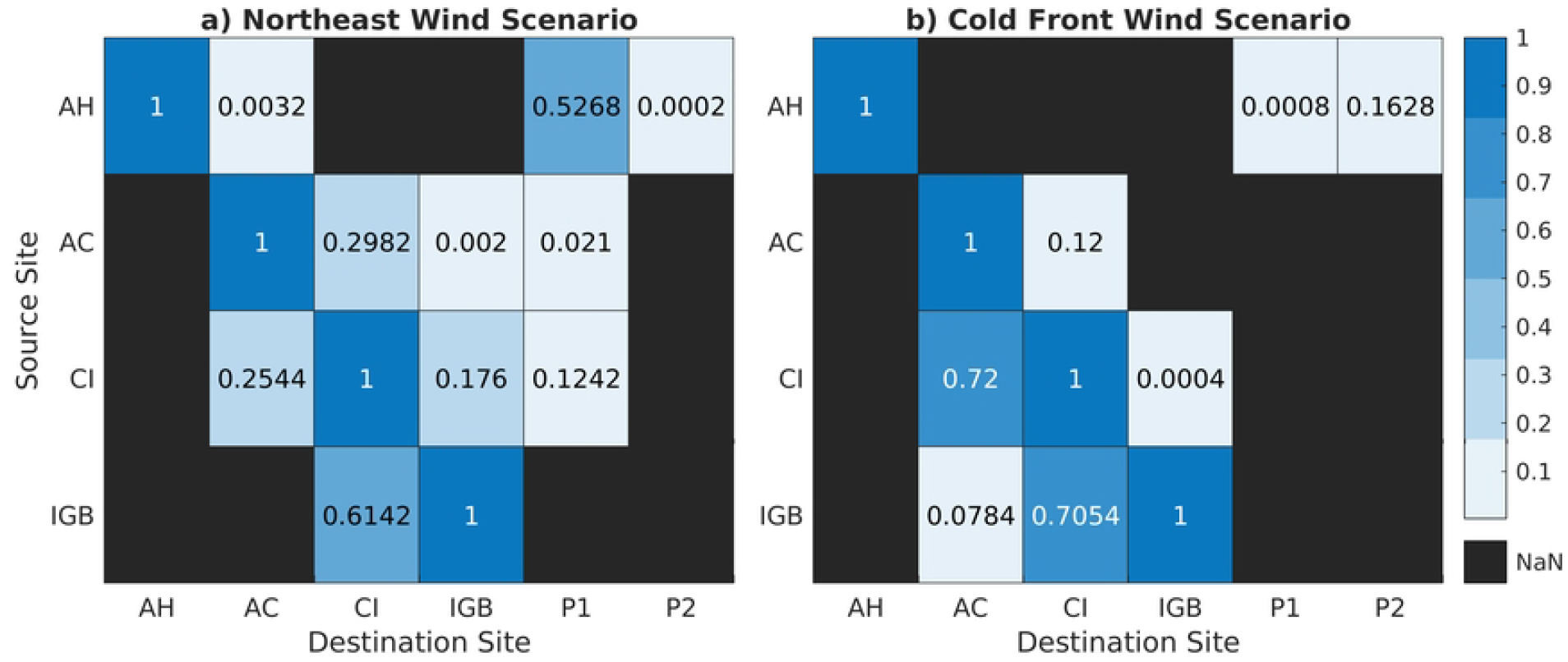
Connectivity matrix of larval dispersal.Results for: a) Northeast Wind Scenario, and b) Cold Front Wind Scenario.

In the northeast wind scenario, illustrated in Figure 8a, there is a greater spatial spread of larvae, with the dispersion extending into offshore regions. Notably, the connectivity between the source sites and offshore points such as P1 and P2 is significant, with P1 showing a percentage value exceeding 0.1. This indicates that the northeast winds, through their capacity to generate offshore-directed export submesoscale filaments, are capable of transporting larvae to distant offshore locations. The connectivity values, while lower than those observed in coastal regions, confirm that larvae are reaching oil and gas exploration zones. This offshore transport, facilitated by submesoscale processes and wind-driven currents, underscores the potential for *Tubastraea* spp. larvae to establish populations on offshore infrastructure, thus expanding their range beyond the coastal zone.

In contrast, the cold front wind scenario, shown in Figure 8b, reveals a more constrained dispersal pattern, with larvae predominantly concentrated along the coast. The correlation values between source and destination sites reflect this limited dispersal range, particularly in the connections between the CI and the AC region, as well as between IGB and the CI. Both of these coastal linkages exhibit percentage values exceeding 0.7, indicating a strong retention of larvae within the coastal system. The higher connectivity values suggest that larvae are less likely to be transported offshore in this wind scenario, remaining confined to nearshore regions where coastal currents and local hydrodynamics dominate their movement.

The contrasting dispersal patterns observed in the two wind scenarios highlight the critical role of wind-driven circulation in shaping the submesoscale transport pathways of *Tubastraea* spp. larvae. The implications of these findings extend beyond the immediate ecological concerns related to *Tubastraea* spp. invasion. The presence of export filaments in the northeast wind scenario suggests that coastal development and offshore infrastructure are inherently connected through these dispersal mechanisms. Offshore oil and gas platforms, which serve as artificial substrates for *Tubastraea* spp. larvae, may act as stepping-stone for the species’ spread. This underscores the importance of monitoring larval transport pathways in both coastal and offshore environments, particularly in regions where human activities, such as oil and gas exploration, may inadvertently facilitate the expansion of invasive species.

Although this study considered only two scenarios, in a real-world context and over an extended period, a greater diversity of dispersion patterns can be observed. The animation in S2 (northeast scenario) and S3 (cold front scenario), which presents simulations over a longer timeframe, reveals that larvae can reach areas beyond the initially predicted regions. Additionally, by incorporating more scenarios into future simulations, different dispersion dynamics are likely to emerge, leading to an even broader range of possible trajectories. This underscores the complexity and variability of larval dispersal processes, which are influenced by multiple environmental and oceanographic factors.

## Conclusion

This study provides an examination of the natural dispersion mechanisms of *Tubastraea* spp. larvae along the southeastern coast of Brazil, focusing on the significant role of wind-driven currents and submesoscale oceanic processes. Through the use of high-resolution numerical modeling, we demonstrate how coastal dynamics—especially submesoscale features, such as filaments and eddies—critically influence cross-shelf exchange. These filaments facilitate the transport of larvae from coastal areas to offshore regions, including oil platforms. This work highlights the complex mechanisms by which biological invasions, particularly involving marine species like *Tubastraea* spp., are facilitated by combined oceanographic and human factors.

A key aspect of our findings is the influence of wind patterns on larval dispersion. Northeasterly winds, which are predominant in the region, drive southwestward sub-mesoscale filaments that transport larval transport offshore. These dynamic features enhance the probability of larvae being transported far from their original coastal habitats, increasing the potential for colonization of distant, offshore ecosystems, including artificial structures like oil platforms. In contrast, southwesterly winds, typically associated with the passage of cold fronts, tend to trap the larvae near the coast, advecting them to the northeast. The interaction between wind-driven currents and submesoscale phenomena thus plays a significant role in shaping the patterns of *Tubastraea* spp. larval distribution.

In addition to natural oceanographic processes, this research draws attention to the influence of offshore oil and gas activities as vectors for the spread of invasive species. Our modeling reveals a clear connectivity between coastal *Tubastraea* spp. populations and offshore installations, suggesting that larvae may be transported either directly by ocean currents or indirectly through human activities, such as the movement of support vessels and industrial operations. The potential for invasive species to colonize offshore structures, such as oil rigs, is of particular concern, as these artificial habitats can facilitate the persistence and further spread of *Tubastraea* spp. populations in marine environments far from their point of origin. This underscores the importance of considering both natural and anthropogenic factors in the management of invasive species.

Finally, this study contributes to the broader understanding of how oceanographic processes shape the distribution of invasive species in marine environments. By elucidating the complex interplay between wind-driven currents, submesoscale features, and anthropogenic factors, we provide critical insights that can inform conservation efforts and the sustainable management of marine resources. Future research should continue to confirm the genetic connectivity of the different populations [52], and to investigate the multifaceted interactions between natural and human-induced factors in biological invasions, with an emphasis on enhancing monitoring capabilities and developing more effective mitigation strategies in vulnerable marine regions. The findings of this study are particularly relevant for policymakers, conservationists, and industries seeking to balance economic activities with the protection of marine biodiversity.

## Acknowledgments

The authors acknowledge the support provided by the Brazilian Navy, Instituto de Estudos do Mar Almirante Paulo Moreira - IEAPM in facilitating this research. We also express our appreciation for the collaborative involvement of ANP (Agência Nacional do Petróleo, Gás Natural e Biocombustíveis) and Equinor Brasil. Furthermore, acknowledge the developers of the Regional Ocean Modeling System (ROMS) for their invaluable efforts (https://www.myroms.org).

## Notes

### Competing Interest Statement

The authors have declared no competing interest.

